# Normative cerebral cortical thickness for human visual areas

**DOI:** 10.1101/676726

**Authors:** Ivan Alvarez, Andrew J. Parker, Holly Bridge

## Abstract

1

Studies of changes in cerebral neocortical thickness often rely on small control samples for comparison with specific populations with abnormal visual systems. We present a normative dataset for FreeSurfer-derived cortical thickness across 25 human visual areas derived from 960 participants in the Human Connectome Project. Cortical thickness varies systematically across visual areas, in broad agreement with canonical visual system hierarchies in the dorsal and ventral pathways. In addition, cortical thickness estimates show consistent within-subject variability and reliability. Importantly, cortical thickness estimates in visual areas are well described by a normal distribution, making them amenable to direct statistical comparison.

**Highlights:**

- Normative neocortical thickness values for human visual areas measured with FreeSurfer
- A gradient of increasing neocortical thickness with visual area hierarchy
- Consistent within- and between-subject variability in neocortical thickness across visual areas

## 2 Introduction

The cerebral cortex is a thin sheet of neurons and glial cells, of which thickness varies between brain regions, reflecting the total neuronal body count, cell type and packing (Nadarajah and Parnavelas, 2002). The heterogeneous distribution of cortical thickness across the human brain appears early in development and develops asymmetrically into adulthood and old age (Gennatas et al., 2017; Hutton et al., 2009; Lemaitre et al., 2012; Salat et al., 2004; Sowell et al., 2004; Thambisetty et al., 2010). Cortical thickness is highly heritable, but developmentally distinct from cortical surface area, volume, or other gross anatomical features (Panizzon et al., 2009; Rimol et al., 2010; Wierenga et al., 2014; Winkler et al., 2010). Instead, cortical thickness appears to reflect the functional organization of the human cortex, with thinner cortex in primary sensory areas and thicker cortex in ‘association’ regions (Van Essen and Glasser, 2014; Van Essen et al., 2012). This observation has led to an interest in cortical thickness as a potential marker for functional specialization in human brain regions and in the development of disease (Van Essen and Glasser, 2014).

While direct measurement of cortical thickness is not currently possible *in vivo*, indirect quantification has become possible through MRI methods, particularly surface-based cortical ribbon reconstruction (Fischl and Dale, 2000). This approach has seen significant refinement with increased spatial resolution and improved signal-to-noise ratio in structural MRI (Fischl, 2012; Lüsebrink et al., 2013). With current MRI acquisition protocols and analysis tools, cortical thickness estimation shows high test-retest reliability (Han et al., 2006; Iscan et al., 2015; Jovicich et al., 2013; Madan and Kensinger, 2017; Wonderlick et al., 2009), good agreement when validated against histological measurements (Cardinale et al., 2014), and an acceptable underlying error rate for group-level statistical comparison (Greve & Fischl, 2018).

In the human visual system, cortical thickness varies significantly between retinotopically organized areas. The primary visual cortex (V1) stands out for its unusually thin band of cortex in contrast to surrounding cortical territory. Cortical thickness may then serve as a marker segregating ‘lower’ sensory from ‘higher’ associative and integrative regions. In addition, abnormal cortical thickness has been reported in developmental disorders (Bridge et al., 2014; Lv et al., 2008), congenital blindness (Bridge et al., 2009; Jiang et al., 2009; Park et al., 2009; Voss and Zatorre, 2012), and progressive illness affecting the visual cortex (Lehmann et al., 2011).

An outstanding issue with these approaches is that measures of cortical thickness in clinical populations are typically compared against a matched control group of similar size, typically less than 20 participants (e.g. (Anurova et al., 2015; Bridge et al., 2009; Lehmann et al., 2011; Voss and Zatorre, 2012). This common approach potentially underestimates the variability of cortical thickness in the population at large, either by measuring too small a sample, or by biasing the control group towards a particular age, gender, ethnicity or other genotypic or phenotypic descriptor in order to match the clinical sample.

In the interest of providing normative values for cortical thickness that better reflect the inter-individual variability in the population, we present a normative dataset of cortical thickness for human visual areas derived from 960 participants in the Human Connectome Project (HCP). The HCP is a rich resource with high-resolution MRI and state of the art pre-processing routines optimized for obtaining anatomical metrics including cortical thickness. In addition to presenting normative values for 25 visual areas, we show that areas are discriminable based on cortical thickness with clusters along the dorsal and ventral pathways, that inter-subject variability in cortical thickness is stable across visual areas, and that within-subject variability is consistent both within and across visual areas. This resource, publicly available on https://ivanalvarez.github.io/NormativeCorticalThickness, can provide a baseline range of values for healthy adult cortical thickness, for researchers wishing to conduct cortical thickness studies in specific populations.

## 3 Materials and Methods

### 3.1 Participants

Data were provided by the Human Connectome Project (Van Essen et al., 2013) S1200 release (release date 01-03-2017). We selected participants who met the following criteria; (1) a complete 3T MRI structural imaging protocol, (2) data processed with the MSMAll registration algorithm (Glasser et al., 2016), and (3) no monozygotic twin pair also present in the sample. Of the participants who met criteria (1) and (2), 286 individuals had a monozygotic twin who was also present in the database with zygosity confirmed with genetic testing. We therefore excluded 143 participants, retaining one participant from each monozygotic twin pair. All dizygotic twin pairs and individuals who self-reported as twins, but were later not confirmed via genetic testing, were retained in the sample. The final sample consisted of 960 participants. The sample has a narrow age range (22 – 37 years), is gender balanced (1 male : 1.17 female) and is drawn from a population with varied demographic, phenotypic and genotypic backgrounds, making it ideal as a baseline for a young adult population more broadly.

### 3.2 Imaging data

MRI data were acquired and pre-processed by the HCP consortium (Van Essen et al., 2013). Structural T1-weighted and T2-weighted images were acquired on a custom Siemens Skyra 3T scanner with sequence parameters optimized for cortical surface reconstruction (Glasser and Van Essen, 2011). In brief, T1-weighted images were acquired with a 3D MPRAGE sequence at 0.7 mm isotropic resolution, and T2-weighted images were acquired with a variable flip angle turbo spin-echo at 0.7 mm isotropic resolution, in addition to B_0_ field maps acquired at 2mm isotropic resolution. Images were pre-processed to correct for distortions introduced by gradient non-linearities, remove readout distortions, correct for bias field distortions and align the images to the MNI space template (Glasser et al., 2013; Goldsmith, 2006; Jenkinson et al., 2002; 2012; Ugurbil et al., 2013). Next, both T1-weigthed and T1-weighted images were used to segment the cortical gray and white matter, and a cortical surface reconstruction generated with FreeSurfer (Dale et al., 1999; Fischl, 2012; Fischl et al., 1999). Morphometric parameters are derived from the surface reconstruction, with cortical thickness estimated as the geometric distance between the white and grey matter surfaces. Finally, the native-space surface meshes were registered to the *164k_fs_LR* template space (Glasser et al., 2013) using multimodal surface registration (MSMAll), which incorporates both anatomical and functional information to improve registration performance (Glasser et al., 2016; Robinson et al., 2013). Resulting surfaces were visualized with the Connectome Workbench software tools (Marcus et al., 2011; Margulies et al., 2013).

### 3.3 Retinotopic areas atlas

Definitions of cortical visual areas were obtained from probabilistic maps of visual topography in (Wang et al., 2015), a publicly available atlas of retinotopic definitions for 25 visual areas in 53 individuals. While cortical visual areas vary in both size and location between individuals (Schwarzkopf et al., 2010), there is high anatomical consistency, particularly for early visual areas (Benson et al., 2014; 2012). Thus, a probabilistic atlas constructed from multiple individuals is a reasonable tool for localizing visual areas in a large participant sample for which individuated visual areas boundaries are not available.

Probabilistic maps for visual area locations were summarized as maximum probability maps (MPM), where each visually-responsive vertex is assigned membership to a single visual area (selection algorithm described in (Wang et al., 2015). Standard space MPMs were projected in a concatenated two-step registration, first to *fsaverage* space in FreeSurfer, and then to the HCP *164k_fs_LR* template. Resulting MPMs are shown in Figure 1. Individual visual area definitions were subsequently used to sample cortical thickness metrics for each HCP subject.

**Figure 1.**
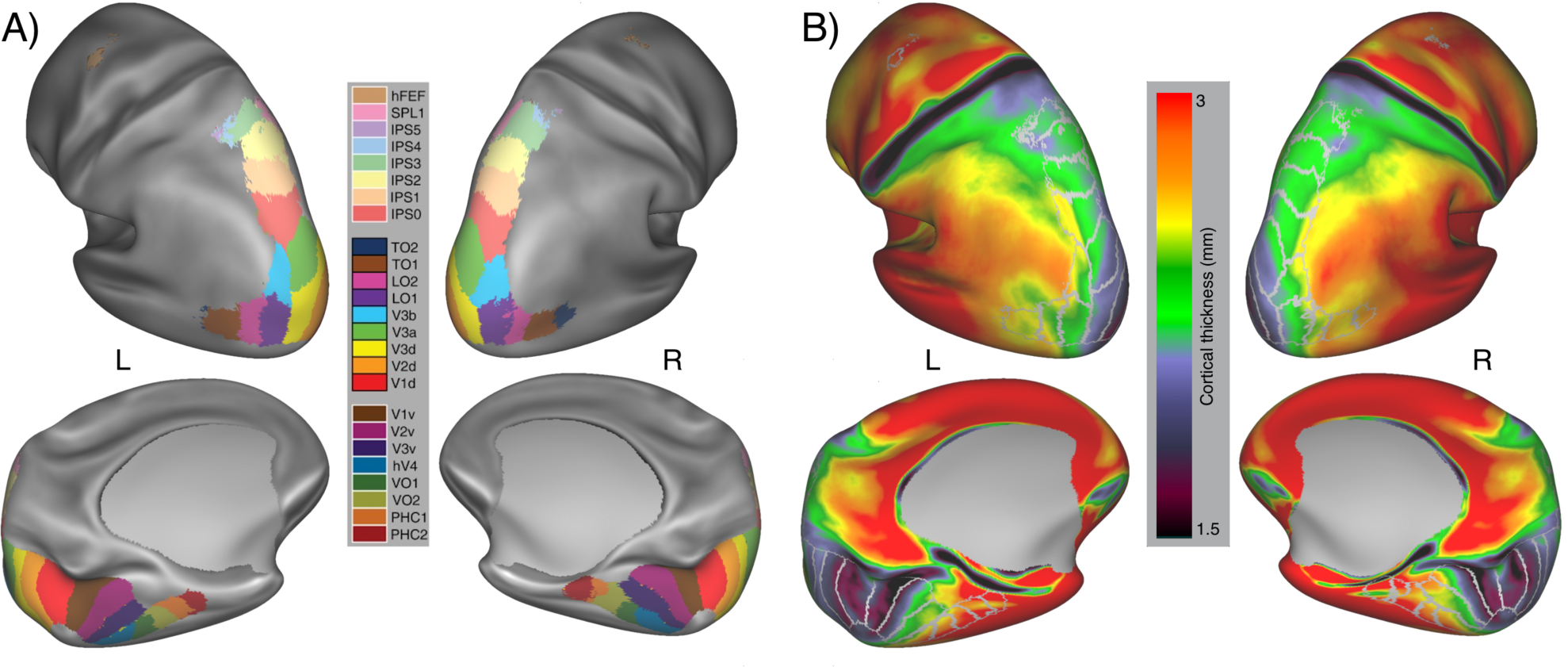
(A) Inflated normalized cortical surface showing the location of visual areas according to their maximum intensity projections in the (Wang et al., 2015) retinotopic atlas, and (B) between-subject mean curvature-corrected cortical thickness for 960 subjects of the HCP dataset. A positive gradient of increasing cortical thickness aligns with visual area cortical hierarchy along both ventral and dorsal directions.

### 3.4 Experimental design and statistical analysis

Surface-based measures of curvature-corrected cortical thickness (corrThickness_MSMAll) were extracted for each subject in 25 regions of interest covering human visual areas that display retinotopic organization. Visual areas were defined by their maximum intensity projection in the (Wang et al., 2015) atlas, and common vertices were assigned to the label with the highest probability. For each area, the mean cortical thickness across vertices was calculated for each subject across both hemispheres, and the standard deviation of the mean taken as a measure of within-subject variability.

Correlation analysis was conducted by taking an individual subject’s mean cortical thickness in each visual area, and correlating across subjects, yielding a correlation matrix of visual area pairs. The correlation matrix was subsequently clustered using a agglomerative hierarchical cluster tree procedure (Bar-Joseph et al., 2001). Statistical analyses were carried out in MATLAB (v9.2, Mathworks Inc., Natick, MA, USA) and R (v3.5.1, R Foundation for Statistical Computing, Vienna, Austria).

The group distributions of mean cortical thickness were fitted with a one-dimensional Gaussian model to obtain standardized population parameters for each visual area. Deviations from the normal distribution were assessed with the Anderson-Darling test, with FDR correction for multiple comparisons. The model goodness of fit was assessed with the coefficient of determination (R^2^) sampled in 100 bins across subjects. The effects of demographic factors on mean cortical thickness were assess with a mixed-factorial ANOVA (3 factors), with age and gender as between-subject factors and visual area as the within-subject independent factor.

Within-subject variability in the cortical thickness was assessed by subjecting it to a ANCOVA, with visual area as the within-subject factor, subject identity as the between-subject factor, while controlling for mean cortical thickness. Reliability of the within-subject mean cortical thickness estimate was assessed with a leave-*p*-out resampling procedure (Celisse and Robin, 2008). In a given visual area, 90% of vertices were drawn with no replacement and averaged to create one sample. For each subject 1,000 samples were obtained, and the span of the 95% confidence interval over the resampled means was then taken as the reliability estimator. Hemispheric asymmetric in cortical thickness was assessed with the two-way, single score interclass correlation coefficient (ICC) (McGraw and Wong, 1996). Within-subject agreement was obtained by comparing left and right hemisphere cortical thickness within subjects. Between-subject agreement was obtained by comparing the left (and right) hemisphere cortical thickness of each subject against the matching hemisphere from every other subject. In addition, a hemispheric bias metric was calculated by subtracting the mean cortical thickness, in mm, for the left hemisphere against the right hemisphere, in each subject, in each visual area. Hemispheric biases were assessed with a series of one-sample t-tests to ascertain if the mean bias across participants originated from a distribution with a mean of zero. FDR correction for multiple comparisons was applied.

## 4 Results

### 4.1 Cortical thickness varies systematically across human visual areas

At the group level, cortical visual areas display significant variability in cortical thickness (Figure 1). In agreement with previous reports of a gradient of cortical thickness along the visual area hierarchy, we observed low cortical thickness estimates in early visual areas V1, V2 and V3 and increasingly larger estimates across the dorsal, ventral and lateral visual areas.

However, it is unclear whether the gradient increase is a feature present at the level of individual subjects, or an emergent feature of group averaging. To answer this question, we conducted a correlation analysis comparing the within-subject mean cortical thickness in one visual area with the within-subject mean cortical thickness in all other visual areas. The resulting Pearson’s correlation coefficients were then grouped using a agglomerative hierarchical cluster tree procedure (Bar-Joseph et al., 2001), with results shown in Figure 2. In agreement with the previous observations, two major clusters of similarity emerge; one in the ventral visual maps, encompassing V2v, V3v, hV4, VO1, VO2, and one in the dorsal visual maps, encompassing V2d, V3d, V3A, V3B, LO1, LO2, TO1 and TO2. Note that V1v and V1d are most similar to each other, with the calcarine sulcus forming the core of small cortical thickness from which the ventral and dorsal clusters span. In addition, a fourth cluster is formed by the higher dorsal-parietal maps IPS0, IPS1, IPS2, IPS3, IPS4, IPS5, SPL1, hFEF, being most similar to each other, and the parahippocampal maps PHC1 and PHC2 form a separate group.

**Figure 2.**
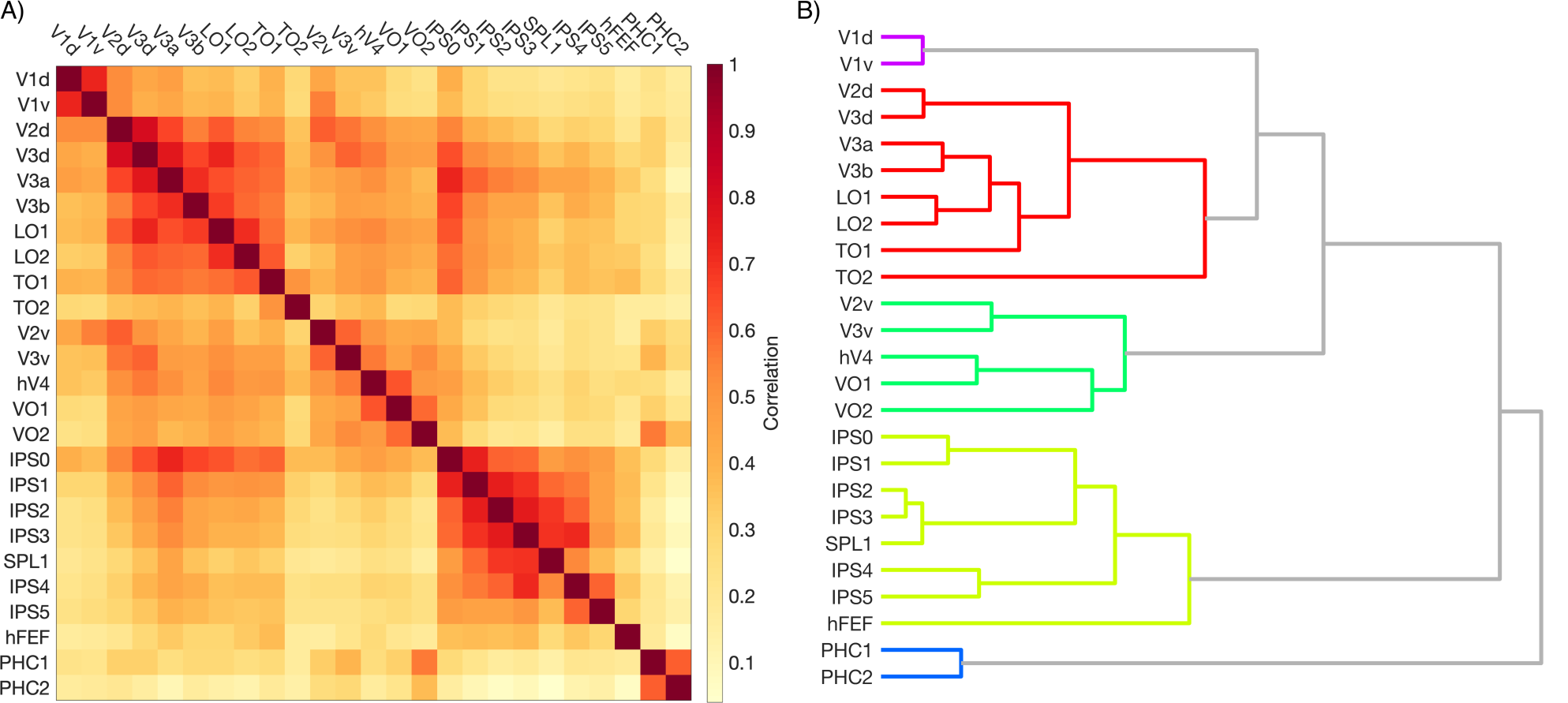
(A) Pearson’s correlation matrix of mean curvature-corrected cortical thickness across 960 HCP subjects in 25 visual areas. (B) Hierarchical cluster tree of correlation coefficients (fully-connected, clustering threshold = 50^th^ percentile linkage height).

### 4.2 Between-subject variability in visual area cortical thickness

Establishing a baseline for between-subject variability in cortical thickness is critical for statistical assessment in small-sampled studies. The large sample size of the HCP dataset allows us to estimate the inter-individual variability in cortical thickness across an adult population with varied demographic backgrounds, as a benchmark for studies of cortical thickness. Surface-corrected mean cortical thickness estimates for each subject, at each visual area across both hemispheres, are shown in Figure 3A. Here, we observe the group variability in cortical thickness is approximately normally distributed in all regions of interest (see Appendix A for quantile-quantile plots). Therefore, we summarize the group distribution as a one-dimensional Gaussian function, also shown in Figure 3B.

**Figure 3.**
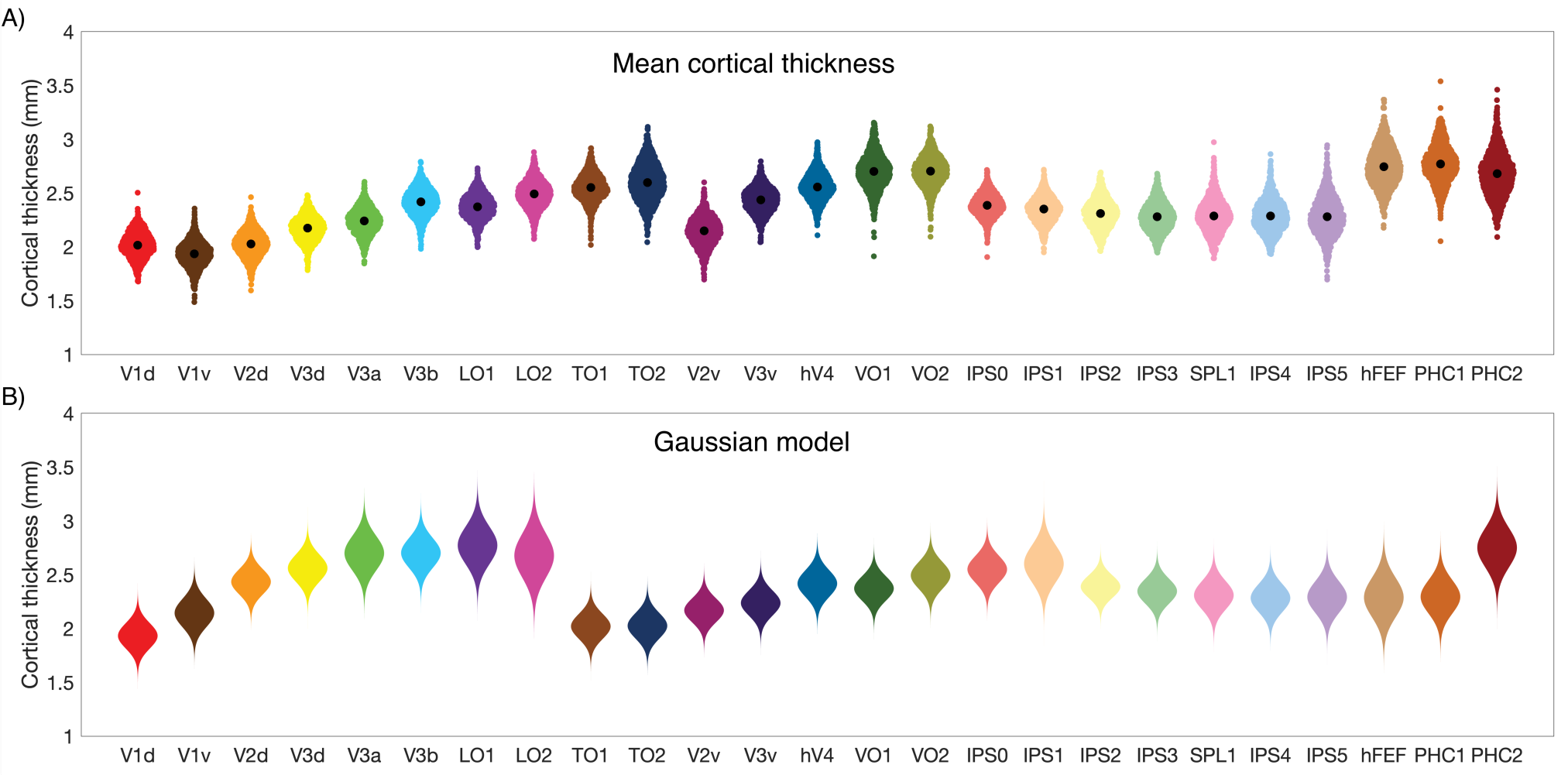
(A) Density plot of mean curvature-corrected cortical thickness across subjects for each regions of interest. Each point, represents the mean cortical thickness for a single subject across both hemispheres, with the center-of-mass at each visual area indicating the group mean. (B) Gaussian distribution model of group cortical thickness for each visual area. Note the two gradients of increasing cortical thickness along the ventral and dorsal visual areas.

In order to assess if cortical thickness estimates are adequately described by the normal distribution, we applied the Anderson-Darling test at each visual area. Twenty-one regions of interest conformed to the normal distribution, with areas VO1, TO1, IPS4 and SPL1 showing significant deviations from normality (FDR-corrected *p* < 0.05, see Appendix B). A two-parameter Gaussian model was fitted, estimating mean cortical thickness peak and dispersion. Performance of the Gaussian model was assessed with the coefficient of determination (R^2^), calculated over 100 equally spaced bins for each visual area (see Table 1). The model explained ≥82% of variance in all regions of interest, including regions registered as non-normally distributed VO1 (92%), TO1 (91%), IPS4 (87%) and SPL1 (88%).

**Table 1.**
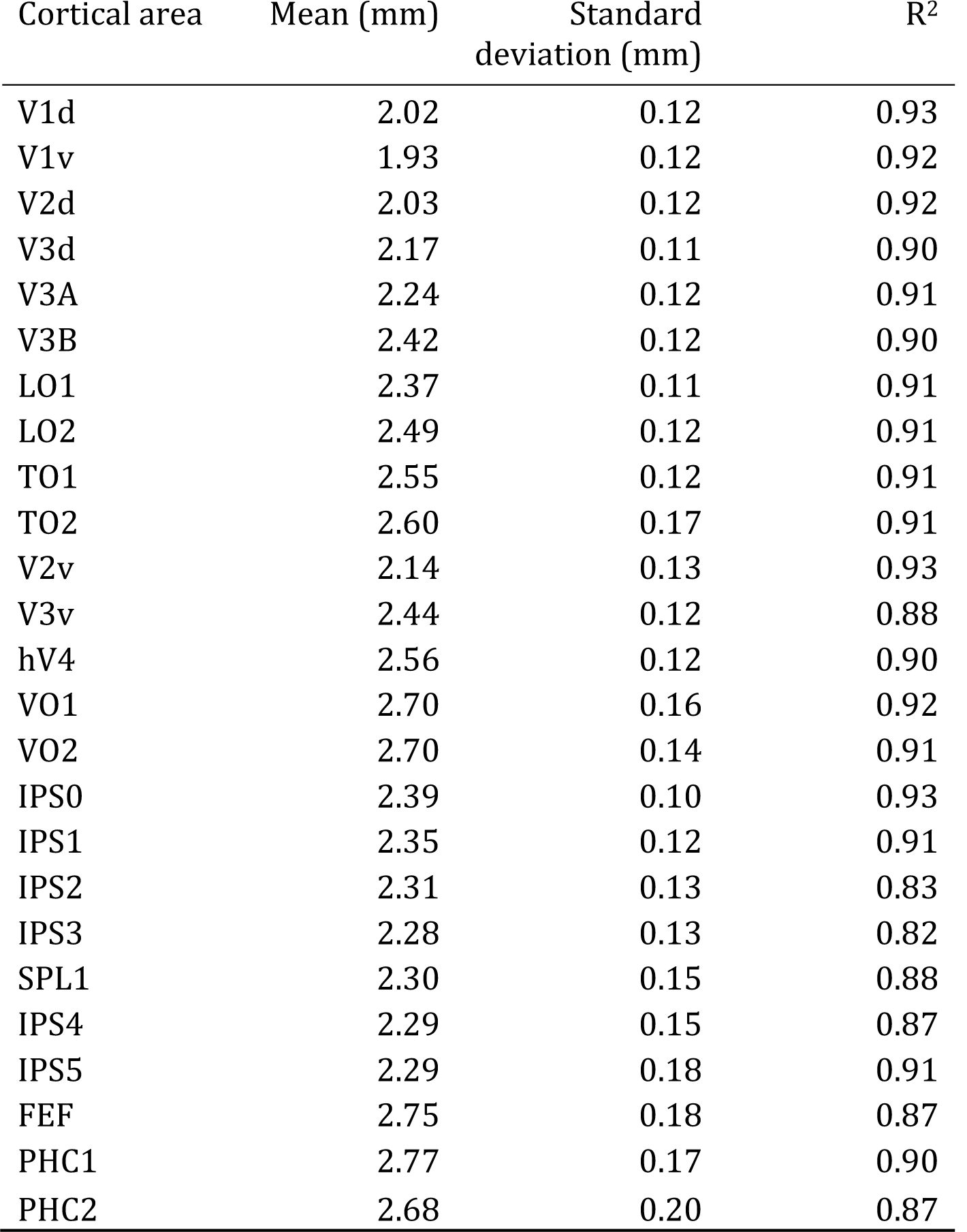
Gaussian model parameters for curvature-corrected cortical thickness across subjects in 25 cortical visual areas. The coefficient of determination (R^2^) was calculated for each area in 100 bins as a measure of goodness of fit. Mean and standard deviation parameters are defined in millimeters of cortical thickness.

### 4.3 Inter-hemispheric differences in visual area cortical thickness

Hemispheric asymmetries in cortical thickness have been previously reported, principally in the frontal and temporal lobes (Im et al., 2006; Luders et al., 2005; Sowell et al., 2004). To examine these effects in cortical visual areas, we calculated the mean cortical thickness per subject in each hemisphere independently.

We assessed whether the cortical thickness in left and right hemispheres of a given subject are more similar than two matching hemispheres from two unrelated subjects. The absolute agreement in cortical thickness was assessed with the two-way, single score interclass correlation coefficient (ICC) (McGraw and Wong, 1996). First, the within-subject agreement was obtained by comparing left and right hemisphere cortical thickness within subjects. Second, the between-subject agreement was obtained by comparing the left (and right) hemisphere cortical thickness of each subject against the matching hemisphere from every other subject. Across visual areas, the within-subject agreement was good (ICC = 0.78), while the between-subject agreement was poor (ICC = 10^-6^). This result confirms the intuition that cortical thickness in the left and right hemispheres is more similar within an individual than matching hemispheres are across individuals.

Following this, hemispheric bias was calculated as the difference in mean cortical thickness between the left and right hemispheres, with positive bias indicating thicker cortex in the left hemisphere and negative bias indicating thicker cortex in the right hemisphere (Figure 4). Hemispheric bias was assessed with independent one-sample t-tests to test if the distribution of biases across participants originated from a distribution with a mean of zero. Of 25 visual areas sampled, 18 displayed significant hemispheric asymmetry (FDR-corrected *p <* 0.05, see Appendix C). Of those areas, 9 displayed medium to large effect sizes (Cohen’s *d* > 0.3), with areas V3v, PHC1, IPS2, PS3 and IPS3 showing thicker cortex in the left hemisphere and areas PHC2, V3d, V3B, and IPS0 showing thicker cortex in the right hemisphere.

**Figure 4.**
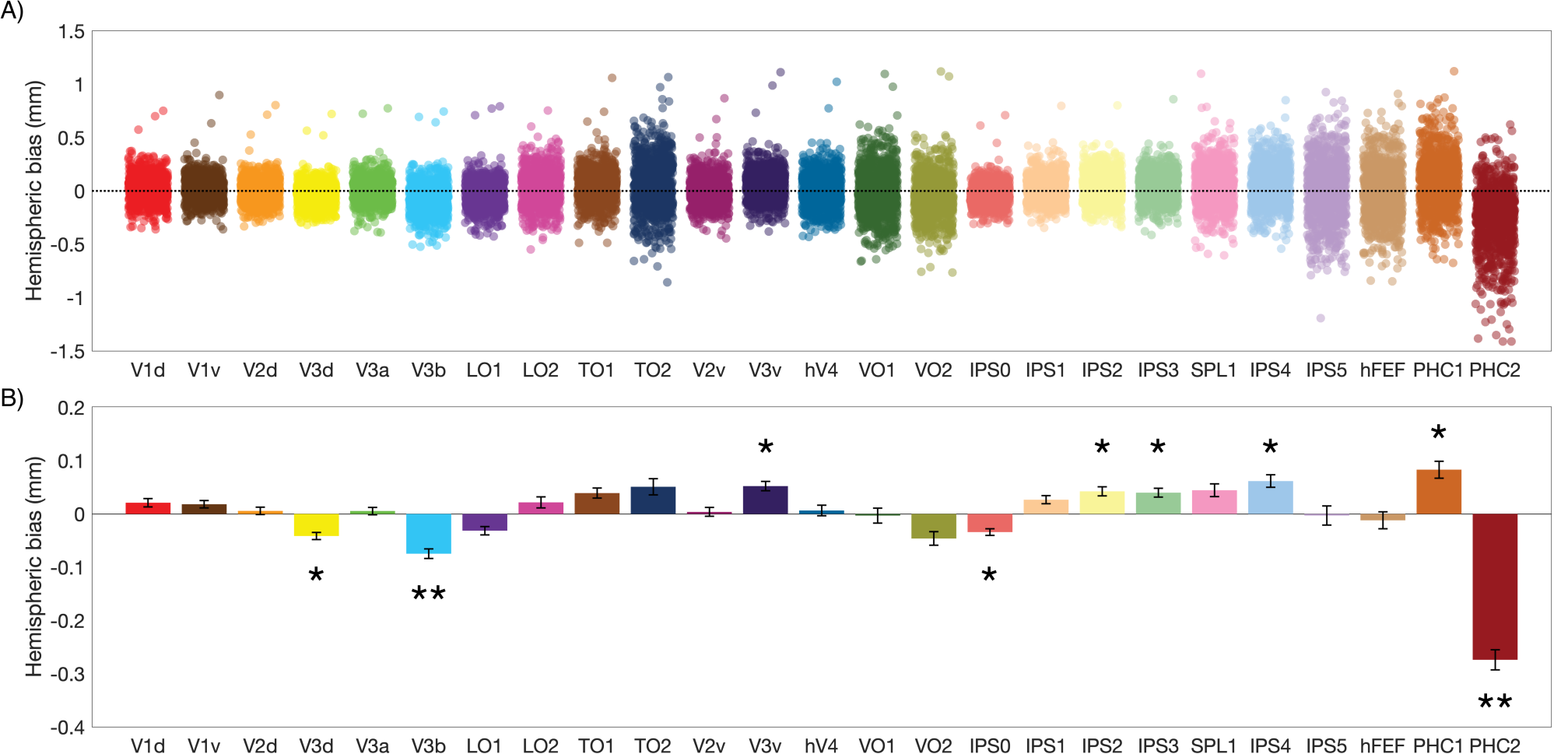
Hemispheric bias for cortical thickness across 25 visual areas. (A) Mean cortical thickness was calculated for the left and right hemispheres independently for each subject, and the difference (left – right) taken as indicator of hemispheric bias. (B) Mean cortical thickness bias across participants and 95% confidence intervals on the mean show areas that significantly deviate from hemispheric symmetry. Areas V3d, PHC1, IPS2, PS3 and IPS3 showed significant bias for the left hemisphere. Areas PHC2, V3d, V3B, and IPS0 showed significant bias for the right hemisphere. * = medium effect size, ** = large effect size.

### 4.4 Age and gender correlates

One factor that predicts cortical thickness is a person’s age. Beyond the initial cortical expansion during childhood development (Sowell et al., 2004), in adults, age is correlated with a decrease in cortical thickness (Salat et al., 2004), and the rate of decrease is non-uniform across the brain (Hutton et al., 2009; Lemaitre et al., 2012; Thambisetty et al., 2010; Wierenga et al., 2014). In addition, previous studies have suggested the relationship between age and cortical thickness may be influenced by gender (Im et al., 2006; Luders et al., 2006; Lv et al., 2010; Wierenga et al., 2014). In order to assess these influences, we examined mean cortical thickness in each region of interest with a mixed ANOVA model, with age in years and self-reported gender as between-subject variables and visual area as the within-subject variable. There was a significant, albeit small, effect of age (*F*(15, 929) = 2.07, *p* = 0.009, η^2^= 0.03), with a mean decrease in cortical thickness of 0.002 mm (± 0.001 SD) per year. This effect is comparable to previous reports of cortical thinning due to normal aging (e.g. (Salat et al., 2004). However, it should be noted the year-on-year slope of cortical thinning falls within the range of measurement error estimates for global cortical thickness (Han et al., 2006; Jovicich et al., 2013; Madan and Kensinger, 2017), as well as within the reliability estimates for this dataset (see Figure 6), and must therefore be interpreted with caution.

**Figure 5.**
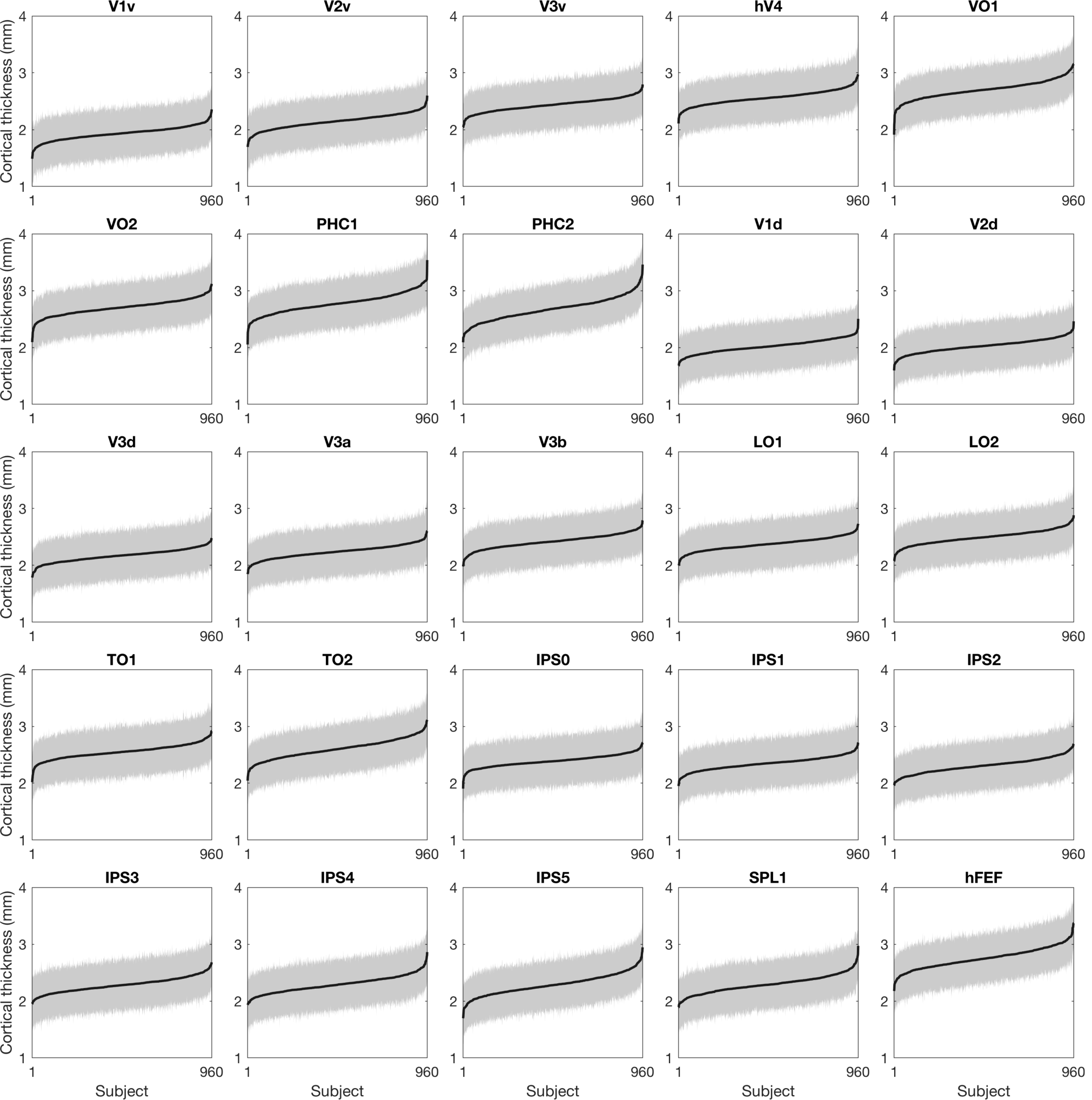
Mean surface-corrected cortical thickness for 960 HCP subjects, across 25 visual areas in rank order. The standard deviation of the within-subject cortical thickness estimate, represented by the grey area, is broadly consistent within each visual area. Mean cortical thickness in black, standard deviation in grey.

**Figure 6.**
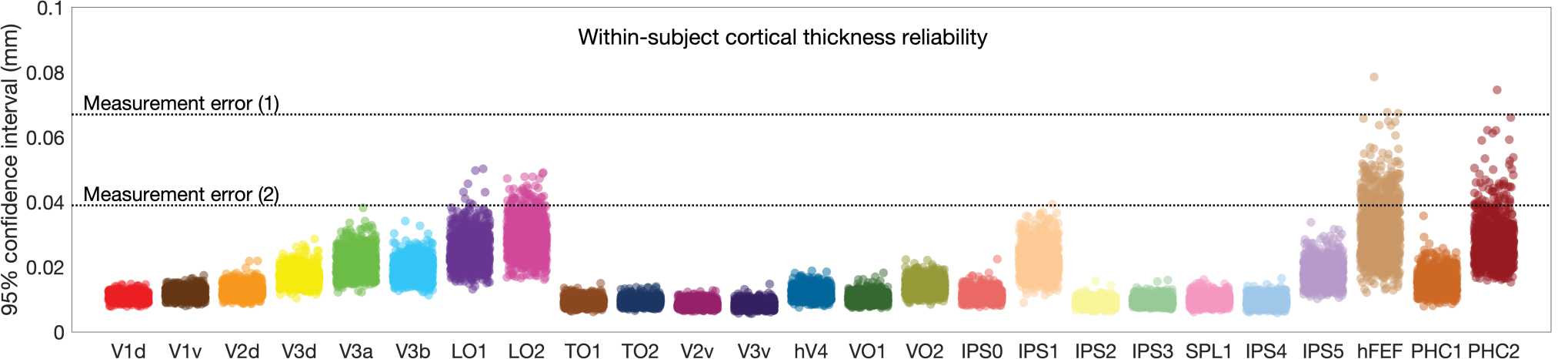
In order to estimate within-subject reliability of mean surface-corrected cortical thickness estimates, a leave-*p*-out resample procedure (1,000 samples, 90% sample size) was conducted, and the 95% confidence interval taken as reliability estimator. One point is displayed per subject. No individual error estimate exceeded 0.1 mm. Dotted lines indicates cortical thickness measurement error as estimated in test-retest studies. Measurement error (1) (Jovicich et al., 2013), measurement error (2) (Han et al., 2006; Madan and Kensinger, 2017).

We detected no effect of gender (*F*(1,929) = 0.12, *p* = 0.729, η^2^= 10^-3^), and no interaction between age and gender on cortical thickness (*F*(14,929) = 1.27, *p* = 0.219, η^2^= 0.02). Summary plots of age and gender effects shown in Appendices D and E, respectively.

### 4.5 Within-subject variability in visual area cortical thickness

The estimation of cortical thickness in a cortical area typically involves assessing the distance between white matter and cerebrospinal fluid at each vertex of the cortical surface. While the individual vertex estimates are unreliable in isolation, over the extent of a region of cortex they form a reliable indicator of the average cortical thickness for that region. However, the within-subject variance for the same cortical region is an informative metric, as it allows us to assess the reliability of the average estimate. Here, we assess the within-subject variability of cortical thickness estimates within each visual region.

We wish to address two questions; first, is within-subject variability comparable across subjects? Second, are the individual mean cortical thickness estimates reliable?

To answer the first question, we looked at the standard deviation of cortical thickness within subjects. For a given visual area, each vertex contains an independent estimate of cortical thickness for that region, and therefore variability in the estimate is observable within areas, on an individual basis. Figure 5 shows the standard deviation of vertex-wise cortical thickness on a subject-by-subject basis. The standard deviation of the within-subject estimate was also assessed with a ANCOVA model, introducing subject identity as the between-subject variable, visual area as a within-subject variable while controlling for mean cortical thickness. No main effect of subject was found (*F*(1, 934) = 0.11, *p* = 0.7417, partial η^2^= 10^-6^), indicating the variance in the cortical thickness estimate is consistent across the HCP population. A significant main effect was observed for visual area (*F*(24, 934) = 543.33, *p* = 10^-16^, partial η^2^= 0.35). This highlights that some regions, particularly the parahippocampal areas PHC1 and PHC2, contain more variability in cortical thickness, either due to true differences in cortical thickness within the cortical areas, or due to unreliability in measurement at this anatomical locus. Finally, no significant interaction effect between visual area and subject was detected (*F*(24, 934) = 0.61, *p* = 0.929, partial η^2^= 10^-3^), showing that despite variability between visual regions, cortical thickness variance is consistent across subjects.

The second question addresses the issue of reliability in the subject-level outcome measure for each visual area, the mean cortical thickness. If a small number of outlier values strongly skews the mean cortical thickness, the outcome measure becomes unreliable. In order to assess the effect of outliers, a leave-*p*-out resampling procedure (Celisse and Robin, 2008) was used where for each visual area, 90% of vertices were drawn without replacement and averaged to create a sample of cortical thickness. For each subject, 1,000 samples were drawn and the span of the 95% confidence interval taken as an estimator of within-subject reliability (Figure 6). Variability in reliability was observed across visual areas, with the highest reliability in early visual areas, as well as dorsal intraparietal areas. Notably, reliability estimates for most visual areas fell below estimates for test-retest error in cortical thickness measurement from three previous studies (Han et al., 2006; Jovicich et al., 2013; Madan and Kensinger, 2017). No individual error estimate exceeded 0.1 mm.

## 5 Discussion

We present a standardized analysis of cerebral cortical thickness in visual areas from 960 subjects made available through the Human Connectome Project.

### 5.1 Cortical thickness and visual system hierarchy

We show that cortical thickness varies systematically across human visual areas, with a positive gradient extending from area V1 into extrastriate areas. A similar pattern of increased cortical thickness with distance from V1 is seen in both the dorsal and ventral pathways, in agreement with previous studies suggesting a link between position in the visual cortical hierarchy and cortical thickness (Wagstyl et al., 2015). When analyzed for similarities in patterns of cortical thickness expansion, dorsal and ventral areas cluster together at the individual subject level, indicating that increasing cortical thickness with visual area hierarchy is not only a feature of averaging cortical thickness metrics over a sample population, but is an identifiable feature in single individuals.

Visual area hierarchy is also reflected in the clustering pattern of cortical thickness correlates between areas, with dorsal and ventral visual areas grouping together. This organization is consistent with the topographical arrangement of retinal representation in visual cortex, where multiple clusters with independent representations of the fovea are found (Buckner and Yeo, 2014; Patel et al., 2014; Wandell et al., 2007). If neighboring visual areas share both common functional architectures and neurodevelopmental trajectories (Cang and Feldheim, 2013), similarities in gross anatomical features such as cortical thickness are likely to arise within visual area clusters.

### 5.2 Population estimates of cortical thickness

Accurate estimation of the natural distribution of cerebral cortical thickness in the population at large is important for studies wishing to compare cortical thickness measured in special populations against a normative baseline. This dataset serves as a baseline for such studies. Between-subject variability in cortical thickness estimates in our sample are well characterized by the normal distribution, with consistent within-subject variability in the sample, and acceptable levels of within-subject reliability in the mean cortical thickness estimate for visual areas.

While the estimates presented here can be considered reliable for early visual areas, those areas at the extremes of visuotopic organization, particularly the intraparietal sulcus and parahippocampal cortex exhibit higher inter-subject variability in their functional localization (Huang et al., 2017; Kolster et al., 2010; Konen and Kastner, 2008; Sereno and Huang, 2014). Indeed, the normal distribution was found to be an insufficient model for areas VO1, TO1, IPS4 and SPL1. In addition to inter-individual variability, extreme dorsal areas IPS4, IPS5 and SPL1 are small, and therefore subject to undersampling in the normalized template space used in this study. Conversely, ventral areas PHC1 and PHC2 lie at a ventromedial locus abutting non-cortical territories, increasing the risk of partial volume inclusion in cortical segmentation. It is unclear if the high variability in cortical thickness estimates observed in PHC1 and PHC2 is a result of measurement error or true anatomical variability within the locus. Overall, cortical thickness values for higher level visual areas should be interpreted with caution.

### 5.3 Relationship to neuronal body count

Cortical thickness, as estimated with MRI methods, reflects a number of factors, including the total number of neurons per unit area, and the ratio of neurons versus non-neuron cells, or neuronal density. In contrast to the findings reported here, stereological studies have identified a negative gradient of neuronal body count along the rostro-caudal axis in non-human primate species (e.g. (Charvet et al., 2014). While neuronal body count influences cortical thickness, the relationship between the two is heavily modulated by neuronal density, which varies between cortical areas (Collins et al., 2010; Herculano-Houzel et al., 2008). The opposite rostro-caudal gradients of human cortical thickness and primate neuronal body count may be indicative of complex intra-area variability in neuronal density, and of potential interest for further study.

### 5.4 Clinical relevance

There are a number of ways in which this type of data can be useful to the interpretation of clinical data. Visual conditions that affect the cortex rather than the retina can be difficult to investigate, particularly when V1 is not affected. Where dysfunction is at the level of V1 or earlier in the visual hierarchy, the resulting visual complaint is usually a loss of visual field. In contrast, when extrastriate visual areas are affected, the subjective reports from sufferers are more confusing, as specific abilities (such as reading) can be affected (Maia da Silva et al., 2017). An example of such a disorder is posterior cortical atrophy, often referred to as the visual variant of Alzheimer’s disease (Crutch et al., 2012). There is clear evidence for cortical thinning, which affects extrastriate areas before V1, although most data are acquired after a diagnosis has been made (Millington et al., 2017). Subtle changes specifically in the pattern of cortical thickness, such as a greater reduction in dorsal areas compared to ventral, or an inversion of the hierarchy, could allow earlier diagnosis of the disease. Thus, even if the populations are not matched for age, the pattern of cortical thickness could still be used. One of the main causes of visual impairment in children in the developed world is now cortical visual impairment (Rahi and Cable, 2003), and the visual profile of this dysfunction is difficult to determine. Like posterior cortical atrophy, the changes in cortical thickness are likely to reflect the specific regions of the visual cortex that are more affected, and therefore could provide additional data to assist with diagnosis.

### 5.5 Compatibility with outside-sample estimates

An important consideration for investigators who wish to use the present study for normative purposes, is whether the values reported here are comparable with other cortical thickness estimates performed outside the context of the HCP. We highlight three relevant considerations; (1) the particular MRI scanner and imaging sequence used, (2) the analysis pipeline implemented and (3) population differences.

First, it is not always possible to match the equipment and sequence parameters across studies, resulting in biases in imaging data leading to differing estimates of cortical thickness. Previous studies have assessed the cross-site, test-retest reliability of FreeSurfer cortical thickness estimates with scans performed on the same individuals, on different MRI systems. Such studies report excellent mean reliability with intraclass correlation coefficients (ICC) ranging between 0.81 and 0.95 (Iscan et al., 2015; Jovicich et al., 2013). In real terms, this corresponds to cortical thickness variability in the range of 0.05 to 0.1mm, depending on the precise cortical area studied. Cross-site reliability values are comparable with reports of within-site, test-rest reliability, with mean ICCs of 0.81 – 0.97 (Liem et al., 2015; Madan and Kensinger, 2017; Tustison et al., 2014; Wonderlick et al., 2009), indicating the effects of scan site are relatively small, making cortical thickness metrics amenable to cross-platform comparisons. Second, the choice of analysis software and analysis pipeline may add additional variability to cortical thickness estimation (Gronenschild et al., 2012; Tustison et al., 2014). However, differences arising at the analysis stage are more easily addressed, either by minimizing discrepancies between the analysis pipeline and that of the normative dataset, or in the case of systematic relationships between pipelines, by applying a scaling factor (Redolfi et al., 2015). Finally, the investigator may wish to examine group differences between a special population, e.g. a clinical group, and the normative data presented here. In such cases, attention must be paid to group differences extraneous to the variable of interest, such as participant motion in the scanner (Reuter et al., 2015), participant age range, and any issues that may selectively affect the accuracy of region of interest localization in one group over the other. Close consideration of these factors will help the investigator determine if statistical comparison with the normative sample presented here is appropriate. Nonetheless, in cases such as posterior cortical atrophy, or cortical visual impairment, the within-subject difference in cortical thickness between visual areas may still be used to indicate those deviating from healthy values.

### 5.6 Conclusions

We present a normative dataset of cerebral cortical thickness for human visual areas in a large sample population in the HCP, with the aim of creating a common baseline for studies of cortical thickness in special populations of interest. Across this large population, thickness varies systematically with cortical visual hierarchy, consistent with the dorsal/ventral organization of the visual system. Moreover, cortical thickness is well described by the normal distribution across visual areas and presents consistent within-subject variability and reliability.

## 6 Appendices

### Appendix A

Quantile-quantile plot of mean curvature-corrected cortical thickness across subjects against the standard normal distribution. Each point represents a single subject across both hemispheres, with the quantile interval represented by a red line. The group-level distribution of mean cortical thickness is consistent with a standard normal distribution across 25 visual areas.

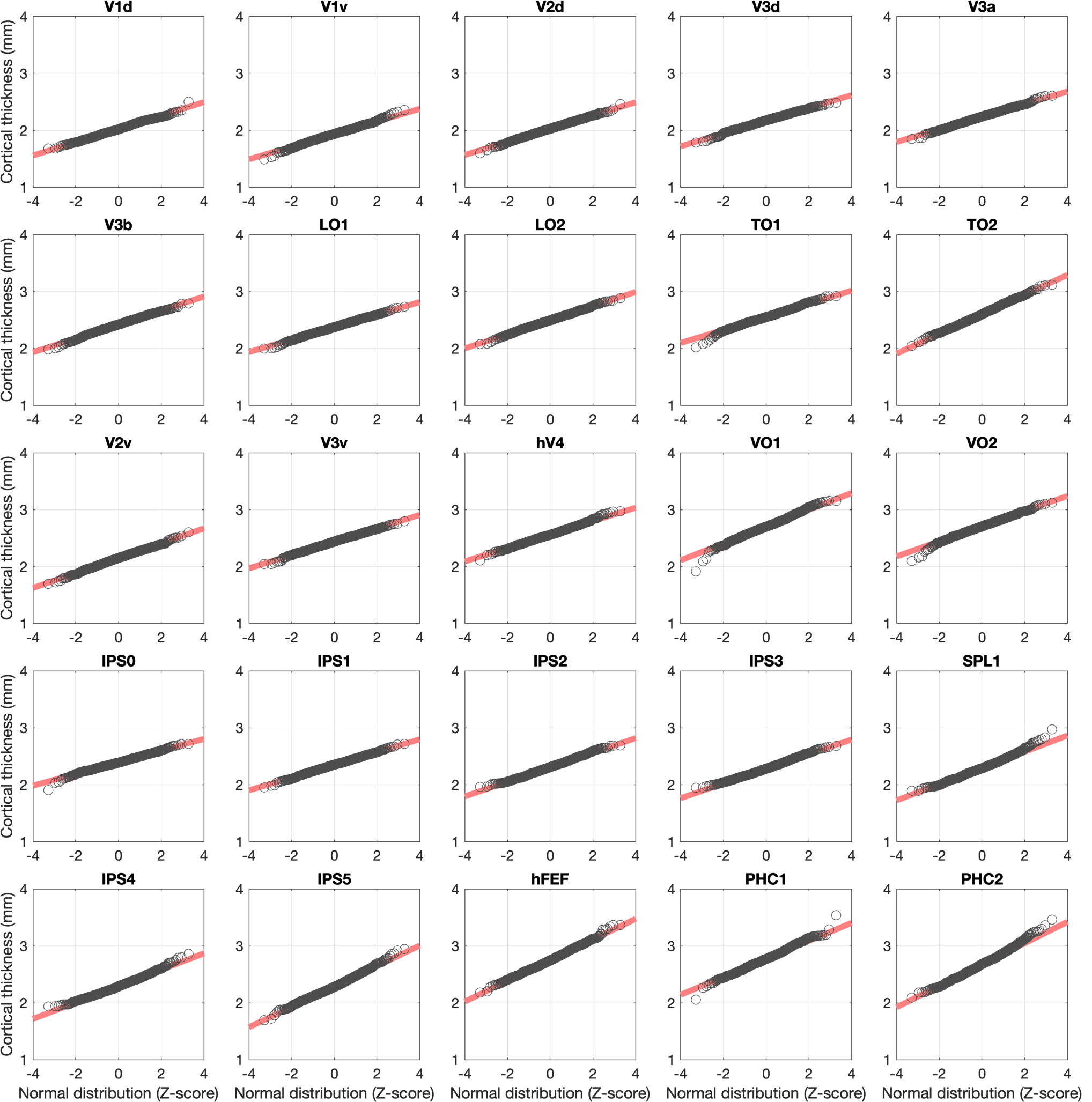

### Appendix B

Anderson-Darling test, assessing whether mean surface-corrected cortical thickness estimates across 960 subjects are consistent with the normal distribution, in 25 visual areas. FDR correction for multiple comparisons applied, **\*** = *p*_*corrected*_ < 0.05.

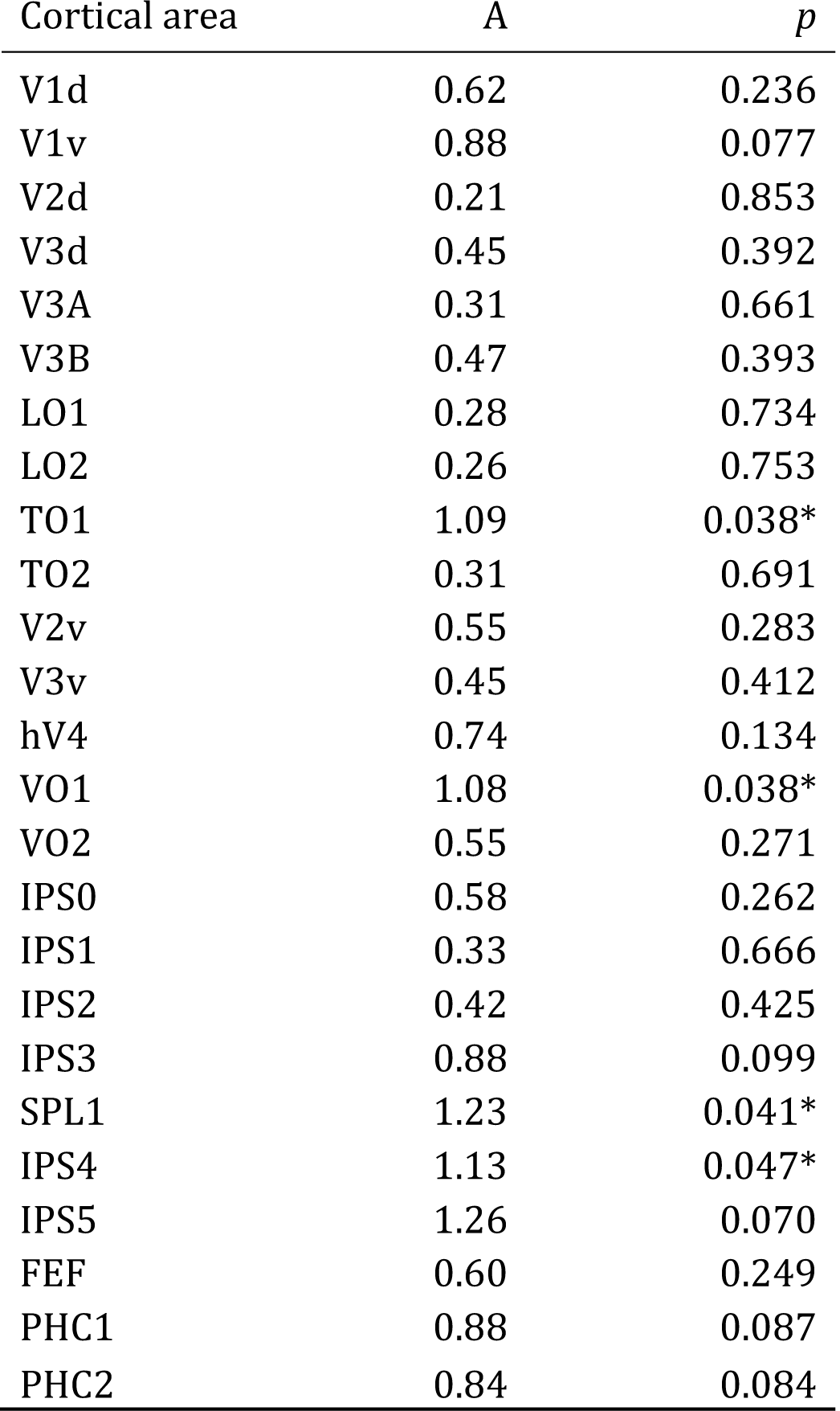

### Appendix C

One-sample t-tests assessing hemispheric bias in surface-corrected cortical thickness (left – right) against a distribution of mean zero in 25 visual areas. Positive hemispheric bias indicates thicker cortex in the left hemisphere, negative hemispheric bias indicates thicker cortex in the right hemisphere. FDR correction for multiple comparison applied, * = medium effect size, ** large effect size.

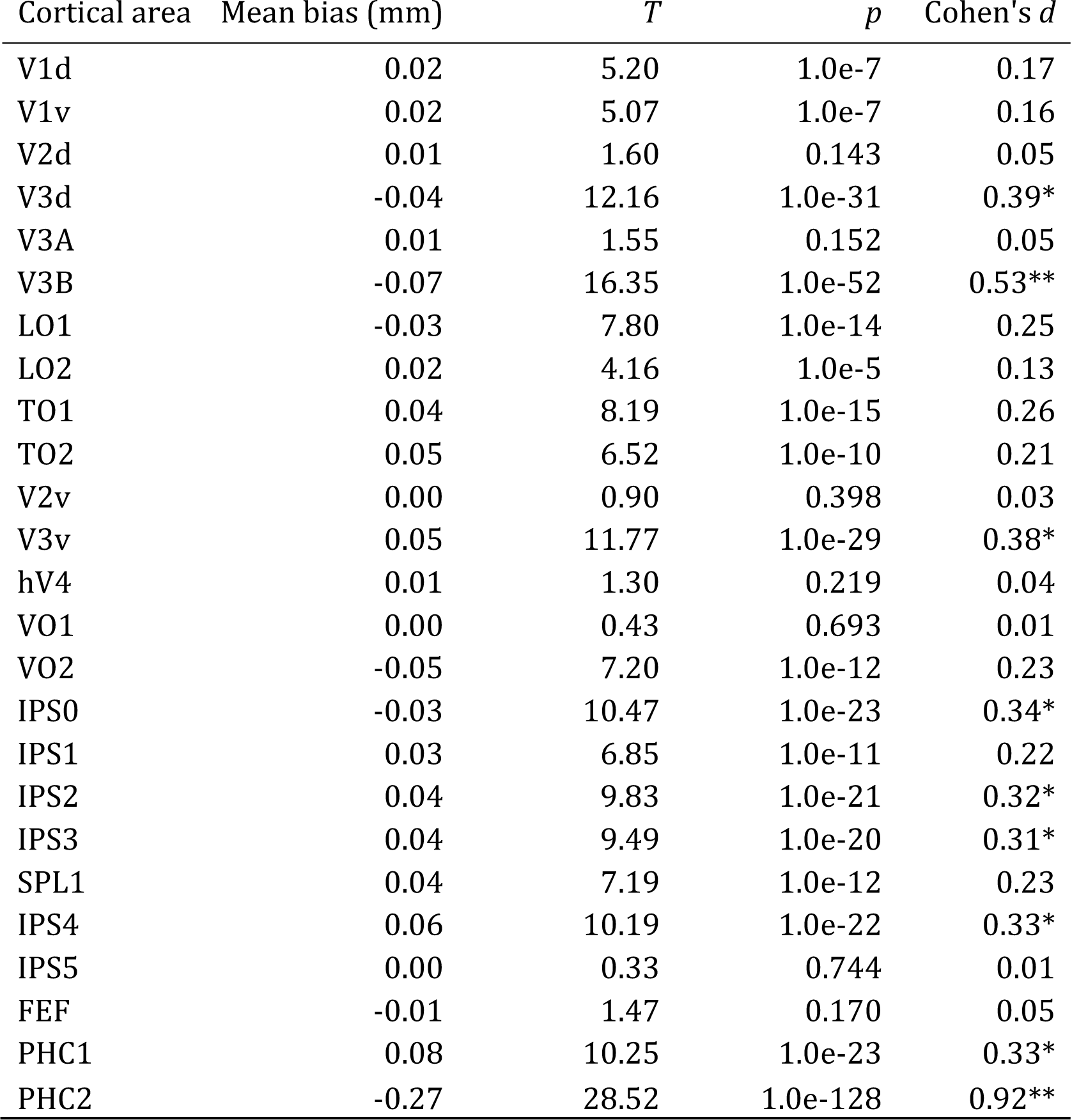

### Appendix D

Relationship between subject age at time of scanning, and mean cortical thickness in 25 visual areas. All cortical areas examined show a negative trend, with cortical thickness decreasing with age at a rate of 0.002 mm (± 0.001 SD) per year. Pearson’s correlation estimates (r) calculated across subjects, best linear fit indicated in red.

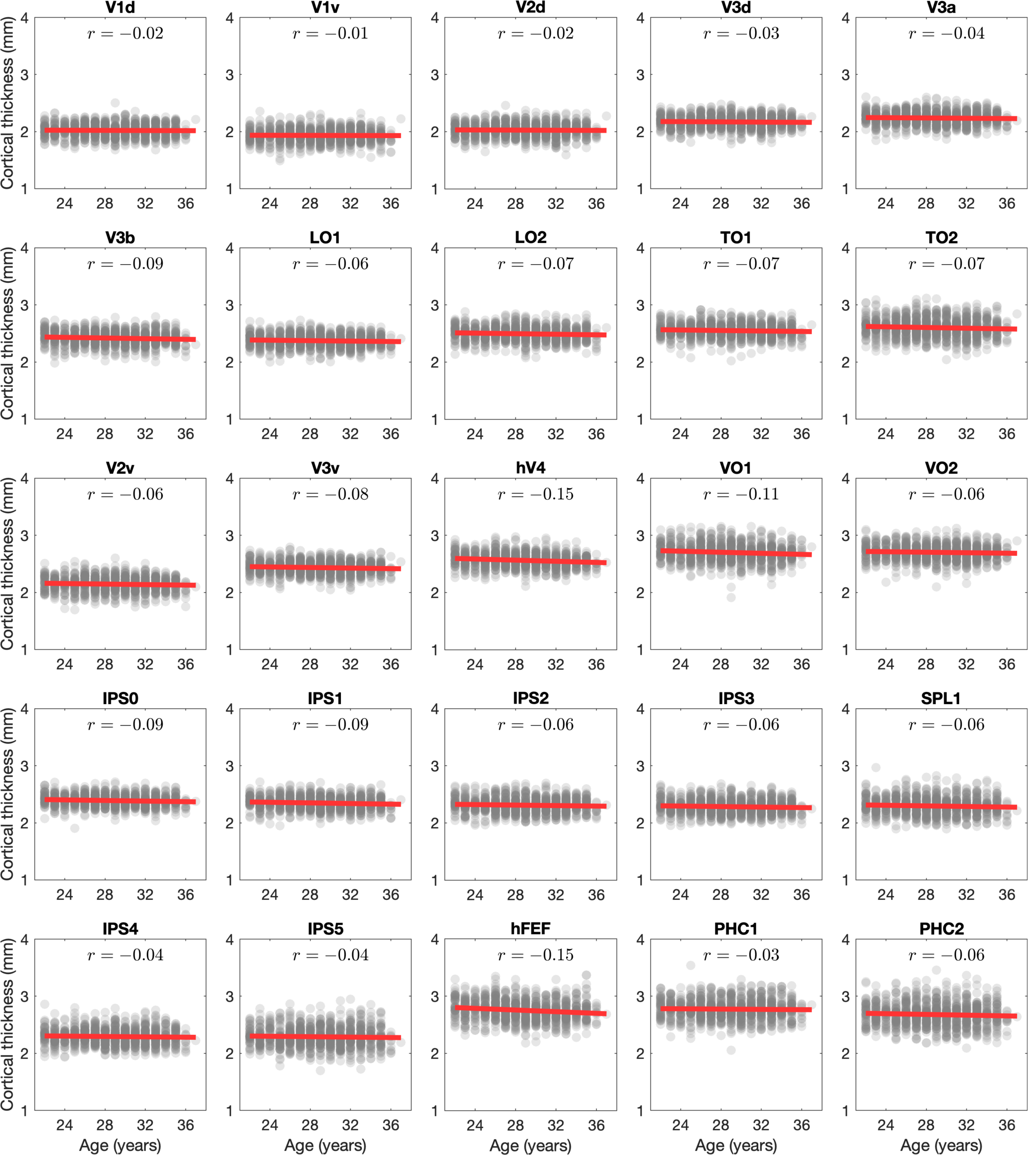

### Appendix E

Relationship between self-reported gender and mean surface-corrected cortical thickness across 25 visual areas. One point displayed per subject. No significant effect of gender, or interaction with age, on cortical thickness was detected.

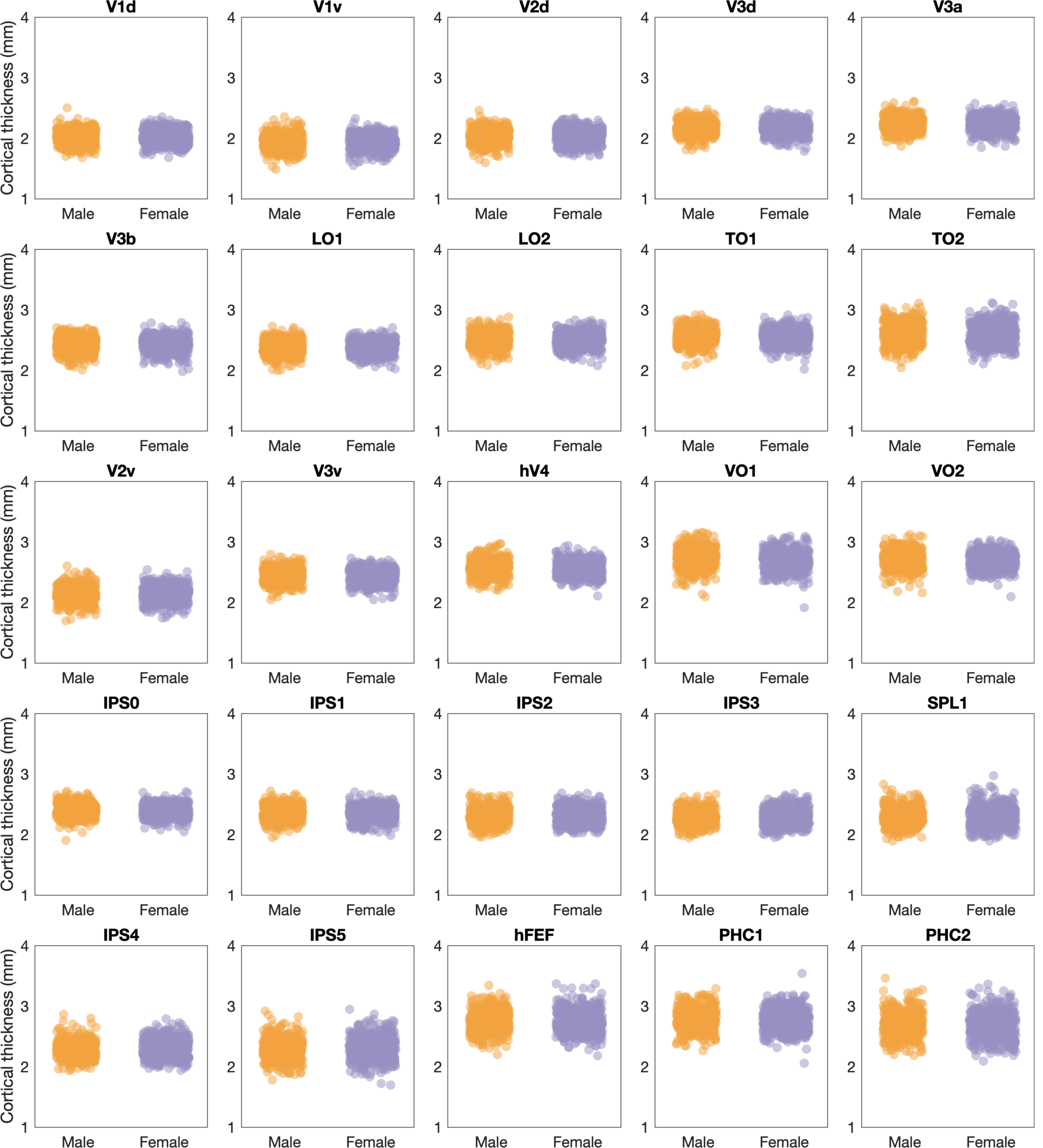

## Abbreviations

(HCP): Human Connectome Project

## Acknowledgements

Data were provided by the Human Connectome Project, WU-Minn Consortium (Principal Investigators: David Van Essen and Kamil Ugurbil; 1U54MH091657) funded by the 16 NIH Institutes and Centers that support the NIH Blueprint for Neuroscience Research; and by the McDonnell Center for Systems Neuroscience at Washington University. This work was supported by the Medical Research Council (MR/K014382/1) and The Royal Society (University Research Fellowship to HB). The Wellcome Centre for Integrative Neuroimaging is supported by core funding from the Wellcome Trust (203139/Z/16/Z).

